# Dietary iron deficiency in the adult mouse increases brain endothelial uptake and blood-brain barrier transport of a high-affinity, anti-transferrin receptor antibody

**DOI:** 10.1101/2024.12.19.629470

**Authors:** Serhii Kostrikov, Kasper Bendix Johnsen, Annette Burkhart, Thomas Lars Andresen, Torben Moos

## Abstract

**Background and objectives:** Brain capillary endothelial cells (BCECs) express transferrin receptor 1 (TfR1) to ensure sufficient iron transport into the brain across the blood-brain barrier (BBB). Our main objective was to examine adult mice subjected to dietary iron deficiency (ID) for possible changes in the content of TfR1 in BCECs and the influence thereof on the uptake and transport of high-affinity anti-transferrin receptor IgG1 antibodies (clone Ri7217).

**Material and methods:** We subjected adult, female mice to dietary ID for 8 weeks. Iron and copper were measured using inductively coupled plasma mass spectrometry (ICP-MS) in various tissues, including total brain and isolated brain capillaries. Possible effects of ID on cerebral angioarchitecture were estimated using 3D confocal microscopy of optically cleared brain samples with endothelium labelled using intravenous injection of wheat germ agglutinin with subsequent machine learning-based segmentation and vascular tracing. TfR1 was quantified using ELISA. Ri7217 antibodies were conjugated with 1 nm nanogold and brain uptake quantified using ICP-MS.

**Results:** ID significantly reduced the iron content in isolated brain capillaries, liver, spleen, kidney, heart and skeletal muscles. ID increased the copper content in the brain. Analysis of cerebral cortex angioarchitecture revealed no changes following dietary ID except for a minor increase in tortuosity of small-caliber vessels. TfR1 protein was unchanged in the total brain and isolated brain capillaries. In contrast, uptake of nanogold-conjugated Ri7217 increased in the total brain, the supernatant fraction of isolated brain capillaries representing the post-vascular compartment, liver, spleen, and dissected retinae.

**Conclusions:** Targeting TfR1 in ID revealed increased uptake and transport across the BBB of Ri7217 antibodies. Possibly the elevated transport of transferrin receptors through BCECs is due to the increased trafficking of transferrin receptor-containing vesicles in ID, which appeared to have no major effects on the brain angioarchitecture.

## Introduction

Iron exerts essential functions in the brain like being a co-factor for heme-containing mitochondrial enzymes and iron-dependent enzymes involved in the synthesis of neurotransmitters (Galy et al., 2024). To ensure sufficient iron uptake, cells express transferrin receptors on their luminal membrane, which allows for the uptake of iron-containing transferrin (Teh et al., 2024). Complicating the provision of iron in the brain, the blood-brain barrier (BBB) formed by brain capillary endothelial cells (BCECs) prevents the passage of transferrin from blood to the brain, which demands for other mechanisms for iron delivery (Taylor and Morgan, 1990). Consequently, BCECs, quite in opposition to capillary endothelial cells elsewhere in the body, have progressed to express transferrin receptor 1 (TfR1) that preferentially acquiesce circulatory iron-transferrin by means of receptor-mediated endocytosis (Jefferies et al., 1984). Subsequently, iron is detached from transferrin within the BCECs and undergoes further transport into the brain, whereas iron-free apo-transferrin is recycled to blood (Taylor et al., 1991).

In BCECs, the uptake and further transport of iron into the brain is increased in iron deficiency (ID) (Taylor et al., 1991). However, somewhat unexpected, the expression of the TfR1 is not increased in ID at the level of the BBB (Moos et al., 1998; Moos and Morgan, 2001). This is contrasted in the early postnatal brain, where TfR1 in BCECs is clearly elevated (Moos et al., 1998). It appears that higher iron transport across the BBB in ID is ensured by enhanced transfer of TfR1-containing vesicles through BCECs (c.f. Johnsen et al., 2019a), possibly assisted by mobilization of a presumably, residing pool of TfR1 (van Gelder et al., 1995).

The TfR1 of BCECs is amendable for drug delivery approaches using specific targeting monoclonal antibodies (Jefferies et al., 1984; Friden et al., 1991). Initially, targeting approaches were based on strategies using monospecific, monoclonal anti-TfR1 antibodies (Jefferies et al., 1984; Friden et al., 1991; Lee et al., 2000). These approaches were later modified by redesign with lowering of their binding properties, e.g. by lowering their avidity, which allowed for higher uptake and transport across the BBB (Yu et al., 2011; Niewoehner et al., 2014) and improved targeted delivery to the brain of iduronate 2-sulfatase (IDS), a lysosomal enzyme deficient in mucopolysaccharidosis type II (MPS II) (Arguello et al., 2022). Recent years, nonetheless, have seen the commercialization of a high-affinity anti-TfR1, which proved able to enter the brain and exert clinical efficacy also by provision of IDS (Sonoda et al., 2018; Harmatz et al., 2024) suggesting that high-affinity anti-TfR1 in conjunction with IDS is also able to pass the BBB.

In the present study, we were intrigued by the idea of using a model of ID in adult mice to study the uptake and possible transport of a high-affinity anti-TfR1, Ri1727, across the BBB. In contrast to the rat, the mouse has only been subject to studies of iron transport at the BBB in few experimental studies, e.g. Baringer et al. (2022). Inducing dietary ID as verified by significant lowering of tissue iron in several organs, we provide a quantitative measure of the TfR1 in isolated brain capillaries in ID. Our analysis of brain angioarchitecture showed no considerable changes in the brain microvasculature, rendering ID as likely not structurally harmful to the brain vasculature, but able to provide higher transport across the BBB when targeted using high-affinity, anti-TfR1 conjugated with 1 nm nanogold.

## Materials and Methods

### Dietary iron-deficiency

Forty-five BALB/c mouse female mice (Janvier Labs) were purchased at the age of 4 weeks and housed in cages at the Animal Department of Aalborg University under constant temperature and humidity conditions and a 12-hour light/dark cycle with free access to food and water. At the age of P42, equal to six postnatal weeks, twenty-three of the mice were shifted from a normal chow diet containing 178.58 iron mg/kg (C1000, Altromin, DE) to the iron-poor diet with a content of 5.2 mg iron/kg (C1038, Altromin, DE) until examined on P98, equal to 14 postnatal weeks. The methods concerning handling of the mice were performed in accordance with the ARRIVE statement guidelines. The Danish Experimental Animal Inspectorate under the Ministry of Food and Agriculture approved all handling of the mice (permission no. 2018−15−0201−01550; approval date October 25, 2018), and the experiments mentioned below were all performed in accordance with relevant guidelines and regulations (EU directive 2010/63/EU) and relevant local additions to this directive.

### Metal analyses

The concentrations of copper, gold and iron were measured by ICP-MS (Helgudóttir et al., 2024) and expressed as ng/g tissue in freshly dissected tissue. Freshly dissected brains were homogenized to isolate brain capillaries, which were separated from the remaining brain, to reveal two fractions, i.e. a capillary enriched fraction and a post-vascular fraction (Johnsen et al., 2018). The samples were digested in aqua regia overnight at 65 °C followed by dilution in deionized water containing 0.5 ppb iridium (Fluka, Sigma-Aldrich, Brøndby, DK). Immediately before analysis, the samples were diluted in 2 % HCl containing 0.5 ppb iridium, after which they were analyzed on an iCAP Q ICP-MS system (Thermo Scientific, Hvidovre, DK) fitted with an ASX-520 AutoSampler and a Neclar ThermoFlex 2500 chiller. Performance on the instrument was ensured by calibration using TUNE B iCAP Q element mixture (Thermo Scientific, Hvidovre, DK). A standard curve was generated by serial dilution of an analytical grade standard solution to obtain the concentration of copper, gold and iron in samples of brain dissected in three fractions, i.e. total brain, brain capillaries and the postvascular compartment. Other dissected tissues were the liver, heart, kidney, lung, spleen, and skeletal muscles dissected from the thigh, retina, and sclera. For dissection of the eyes, both eyeballs were removed in toto, and the retinas and sclerae were gently dissected using small forceps under a surgical microscope. Copper measured in the sclera and retina is not reported as measures were below detection level. Measurement of the iridium concentration was used as an internal standard to ensure similar analysis of all samples.

### Labeling, visualization and quantitative binding of anti-TfR1 antibodies to the vascular tree in normal and ID mice

#### Vasculature labelling and sample preparation

To achieve vasculature labelling, mice received tail-vein injection of 200 μl AF555-conjugated Wheat Germ Agglutinin (WGA) lectin (1.5 μg/μl in PBS). After 10 minutes of lectin circulation, mice were anaesthetized with 8% sevoflurane and transcardial perfusion was performed. First 30 ml of PBS was perfused, which was followed by a perfusion of 20 ml of 4% methanol-free formaldehyde (Thermo Fisher Scientific, #28908). Brains were dissected and postfixed in formaldehyde solution for 24 hours at -4°. Postfixed brains were cut in at the distance of 1 mm and 3 mm from Bregma. Obtained 2 mm-thick samples underwent serial dehydration in tetrahydrofuran (THF) (Merck, #186562) in MilliQ water as follows: 3 hours in 30% THF, 3 hours in 50% THF, overnight in 70% THF, 3 hours in 90% THF, 3 hours in 100% THF, overnight in 100% THF. All steps were performed at room temperature with gentle agitation of the samples. Refractive index matching was done in ethyl cinnamate (Sigma Aldrich, #112372) for at least 3 hours before imaging.

#### Microscopy

Imaging of optically cleared samples was performed using confocal laser scanning microscope (Zeiss Microscopy, LSM 710) with 0.3 NA 10x objective. For the fluorophore excitation, 561 nm laser was used. The microscope settings parameters were adjusted to acquire the signal intensity just below the saturation limit; with the same purpose, Z-correction was used. Images were taken in the cerebral cortical region knowingly that this might denote a potential limitation of the study that only this region was examined.

#### Vasculature network analysis

For the vasculature network analysis, we used approach developed previously (Kostrikov et al., 2021) with slight modifications. First image datasets underwent blind deconvolution in Amira 6.7 (Thermo Fisher Scientific). Deconvolution was followed by the histogram equalization adopted from Baalousha et al. (2015). Pixel intensities were modified as described by the formula below:

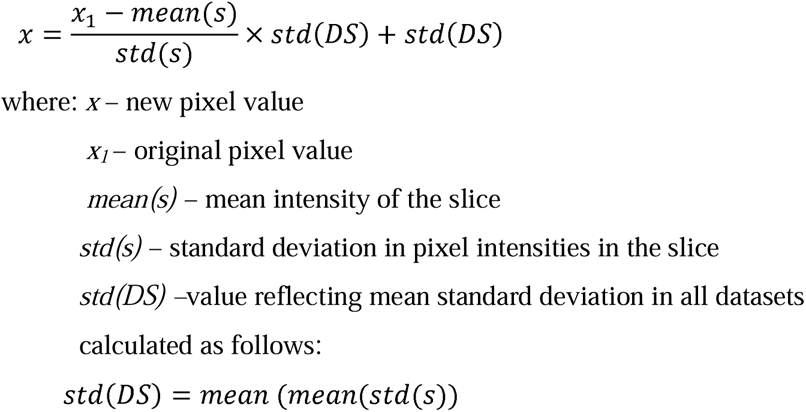

Vasculature of the cortical angioarchitecture in the equalized datasets was segmented using machine learning-based pixel classifier – Zen Intellesis (Zeiss Microscopy). Obtained binary masks were then postprocessed. First 3D Gaussian smoothing was done (σ = 1) followed by thresholding (pixel intensity: 80-255). Hollow vessel lumens were filled using the approach developed by Kostrikov et al. (2021). Thereafter, masks were further smoothed with 2D median filter (radius = 1). All the postprocessing operation were done in FIJI. From obtained images, Euclidean distance map was generated in Amira. After distance transformation, spatial graph of vasculature was generated. The obtained spatial graph was then smoothened in Amira using Smooth Line Set module with smoothing coefficient equal to 0.9, adherence to original data coefficient equal to 0.07 and number of iterations equal to 15.

To maximize the precision of vessel diameter measurements, correction factor was calculated based on the manual measurements of 100 segments. Obtained formula for the correction factor was as follows:

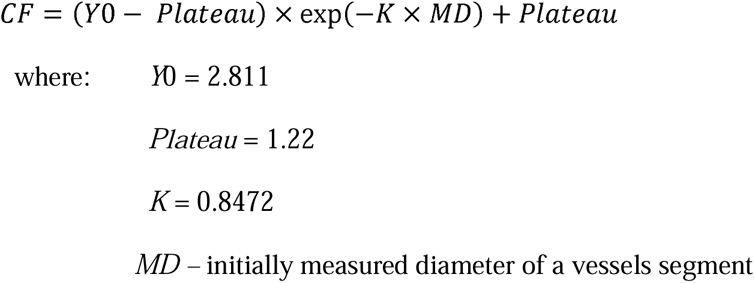

### Labeling of anti-TfR1 antibodies with gold nanoparticles

To estimate the uptake of TfR1 in the brain and peripheral organs, we used the Ri7217 antibody labeled with 1.4 nm gold nanoparticles (Nanoprobes Inc., Yaphank, NY, US) for high-sensitivity detection using ICP-MS. A stock buffer (PBS) of the anti-TfR1 antibody was exchanged to 0.1 M sodium borate buffer with 2 mM EDTA (pH 8). The buffer-exchanged antibodies were then subjected to thiolation using Traut’s reagent (Thermo Scientific, Hvidovre, DK) at a reagent-to-antibody ratio of 10. The solution was allowed to incubate for one hour at room temperature under constant shaking at 500 rpm. The resulting thiolated antibodies were then transferred to an Amicon Ultra spin filter (MW cut-off 30 kDa) (Sigma-Aldrich, Søborg, DK), topped with 5 mL sodium borate buffer, and centrifuged at 4000 g for 20 min at room temperature. The filter-through was discarded, and 5 mL PBS (pH 7.4) was added to the thiolated antibodies. After volume reduction to 50 µL by centrifugation, the thiolated antibodies were transferred to a Protein Lo-Bind Eppendorf tube and stored at 4°C for no more than 30 min. Conjugation of thiolated antibodies to the 1.4 nm gold nanoparticles was then performed using monomaleimide Nanogold Labeling Reagent (Nanoprobes Inc., Yaphank, NY, US) according to the manufacturer’s instructions. In short, the lyophilized monomaleimide nanoparticles were mixed in 1 mL of deionized water and added to 2 mg thiolated antibodies. The air phase was replaced with N_2_, and the solution was incubated at 4°C overnight. The next day, the labeled antibodies were separated from unbound gold nanoparticles using gel filtration chromatography on a Superose 6 column (Sigma-Aldrich, Søborg, DK) and stored at 4°C until use.

### TfR1 analysis

ELISA was performed to measure the amount of TfR1 in brain homogenates and isolated brain capillaries. Freshly dissected brains of non-injected mice fed a normal (n = 6) or ID (n = 6) diet were homogenized to isolate brain capillaries. An initial measure of the protein concentrations was measured using a BCA protein assay kit as previously described (Helgudóttir et al., 2024). Each sample was lysed using 90 µL N-per Neuronal protein extraction buffer supplemented with cOmplete Mini protease inhibitor. The samples were lysed on ice for 10 min and spun down at 14,000 g for 10 min at 4°C and the supernatant collected. Samples were diluted 1:10 in sample buffer and analyzed using sandwich-ELISA for mouse TfR1 according to the manufacturer’s protocol. The absorbance was read at 450 nm using an Enspire Plate Reader from Perkin Elmer, and the concentration calculated using a standard curve normalized to the total protein concentration of each individual sample.

### Uptake of nanogold-conjugated anti-TfR1 antibodies

Mice fed a normal (n = 8) or ID (n = 9) diet were intravenously (iv) injected with nanogold-conjugated anti-TfR1 antibodies dissolved in 0.1 M PBS, pH 7.4 in a dose of 5 * 10^11^ AuNPs/mouse. The animals were monitored for 30 min and then euthanized by transcardial perfusion with 20 mL 0.01 M KPBS (pH 7.4) under isoflurane anesthesia. Brains (one hemisphere) and peripheral organs of interest were dissected, snap-frozen in liquid nitrogen and kept at -80°C until further analysis for the content of Au, Cu and Fe (Johnsen et al., 2018). For estimating the transport across the BBB, one hemisphere was homogenized and subjected to the brain capillary procedure as previously described (Johnsen et al., 2018; Johnsen et al., 2019b). For dissection of the eyes, both eyeballs were removed in toto, and the retinas and sclerae were gently dissected using small forceps under a surgical microscope. The tissue of mice injected with anti-TfR1 antibodies was also used for the displayed analyses of iron and copper described above.

### Statistics

All data were analyzed for outliers using the ROUT test and for equal variances using an F-test. The statistical analyses were two-sided, and a p-value of ≤ 0.05 was considered statistically significant. Normality of the distribution of residuals was estimated using Shapiro-Wilk test. In case of normal distribution of residuals, an unpaired t-test (if variances were equal) or a Welch test (if variances were not equal) was used. If distribution of residuals was not normal, Mann-Whitney test was used. Animal weight was analyzed using a two-way ANOVA with Šídák’s multiple comparisons test. Graphical presentation and statistical testing were created using GraphPad Prism 10.4.0 and data are depicted as mean ± standard deviation (SD) for barplots and mean ± standard error of mean (SEM) for lineplots. N values refer to the number of animals included in the analysis. The details of the specific analysis and n-values are reported in each figure legend.

## Results

The mice were thriving following an 8-week period of dietary ID and followed the same weight curve as seen in the control-fed mice with a weight gain from approximately 19.5 grams to 24 grams (Fig. 1A). The iron content of the total brain was 13.9 ±0.67 µg/g for normal feed mice and 13.6 ±0.08 µg/g for the ID-fed mice (Fig. 2). These values did not vary change significantly. In contrast, a lowering of iron was seen following capillary isolation in both the pellet fraction representing the brain capillaries and the supernatant fraction representing the post-vascular brain compartment (Fig. 2), e.g. in the isolated brain capillaries iron differed from 4 µg/g in control fed mice to 3 µg/g in ID-fed mice (p < 0.001). In extracerebral tissues, iron dramatically lowered in ID-fed mice with significantly lower iron content in dissected samples of the liver (p < 0.0001), spleen (p < 0.0001), skeletal muscle (p < 0.001), heart (p < 0.0001), kidney (p < 0.0001), and sclera (p < 0.05). Examining samples from the same mice revealed that copper essentially remained at constant levels except for a significant increase in dissected samples of the total brain from approximately 3.7 ± 0.07 µg/g to 4 ± 0.2 µg/g (p < 0.005) (Fig. 3). A slight decrease in skeletal muscle content of copper was also seen (p < 0.005).

**Figure 1.**
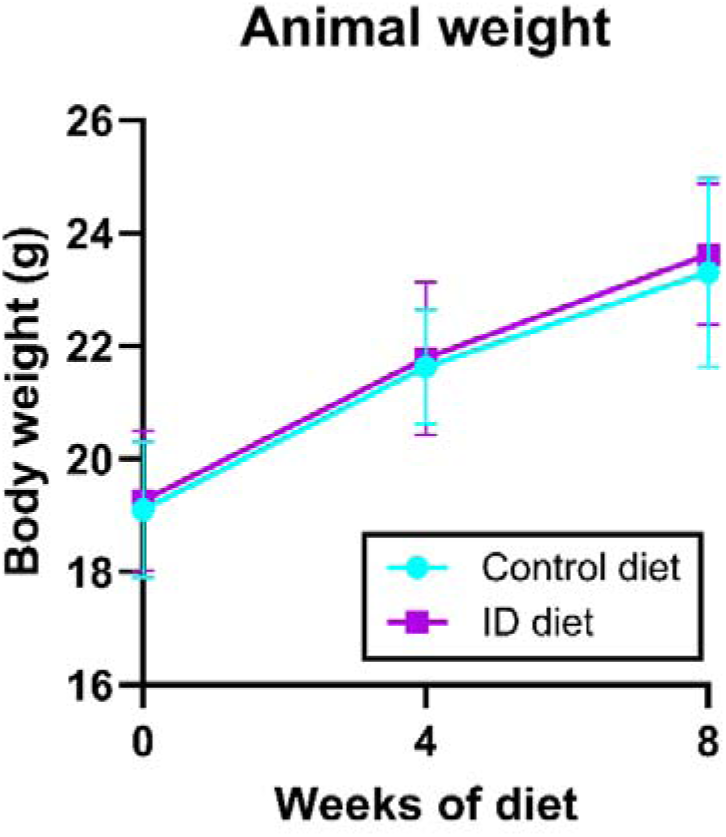
Development of body weight in mice fed a control (turquoise) or ID (magenta) diet. Feeding the mice with the ID diet for 8 weeks allows the mice to follow the same development of the weight curve as seen in the control-fed mice. Data are analyzed using a two-way repeated-measures ANOVA with a Šídák’s multiple comparisons test. There were no significant differences in body weight between the two groups. Data are depicted as mean ± SD (*n* = 19 mice/group).

**Figure 2.**
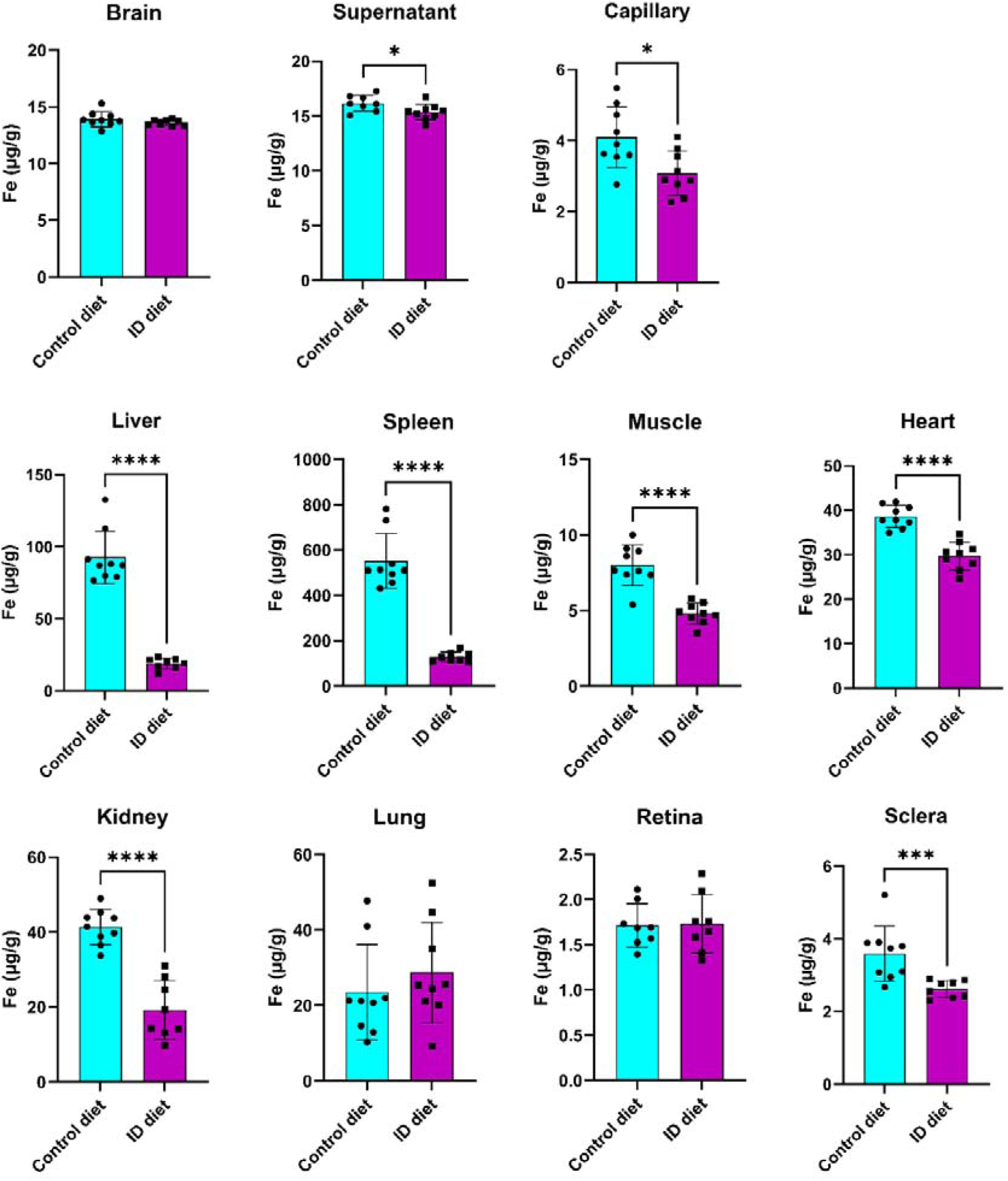
Iron (Fe) content in mice fed a control (turquoise) or ID (magenta) diet. Feeding the mice with the ID diet leads to a significant lowering of Fe in the supernatant and pellet fractions of the brain capillary isolation (p < 0.05), liver (p < 0.0001), spleen (p < 0.0001), skeletal muscle (p < 0.0001), heart (p < 0.0001), kidney (p < 0.0001), and sclera (p < 0.001). Data are analyzed using an unpaired t-test or a Mann-Whitney test (Brain, liver, spleen, and sclera). Data are depicted as mean ± SD (n = 8-9 mice/group).

**Figure 3.**
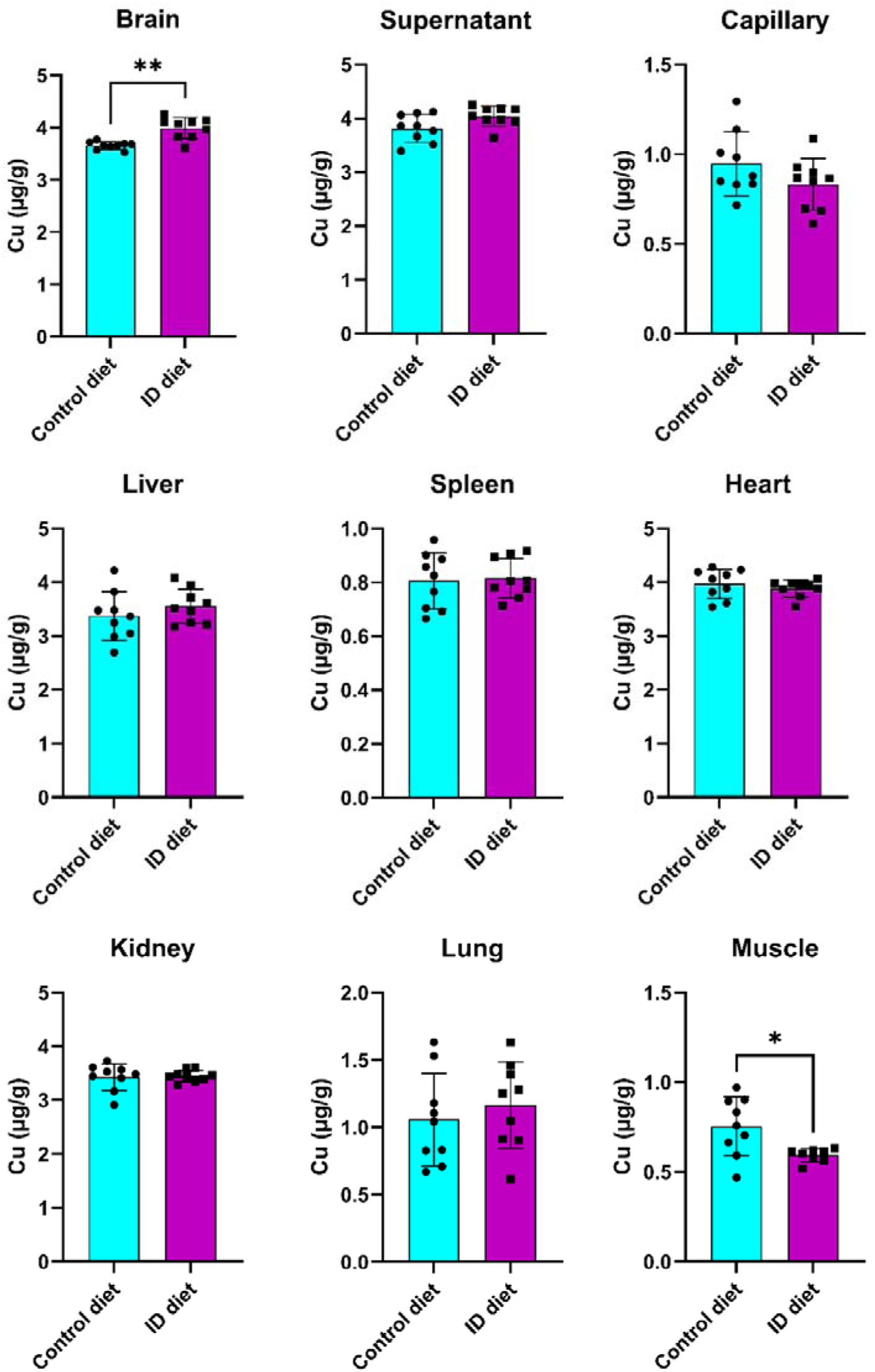
Copper (Cu) content in mice fed a control (turquoise) or ID (magenta) diet. Feeding the mice with the ID diet increases the Cu content in the total brain (p < 0.01) and decreases the Cu content in the skeletal muscle (p < 0.05). The sclera and retinas are not shown due to limitations in sensitivity. Data are analyzed using an unpaired t-test or a Mann-Whitney test (brain, kidney, and muscle). Data are depicted as mean ± SD (n = 8-9 mice/group).

3D analysis of cortical angioarchitecture showed no major changes (Figs. 4A-L). No changes in the ratio between perfused endothelial surface and extravascular space volume (Fig. 4 I), suggest unimpaired capacity of the vasculature to provide gas and nutrients exchange. Vessel segment length was also unchanged (Fig. 4K), suggesting no ongoing angiogenesis. This was also corroborated by no changes in the vessel diameter (Fig. 4J) The mean vessel tortuosity calculated as the ratio between the actual segment length and the chord length (smallest distance between segment start and segment end) for all segments in an image was also not changed; however, the subpopulation analysis showed a small but significant increase (p < 0.05) in the tortuosity of the small-caliber (<4 μm) vessels (Fig. 4L).

**Figure 4.**
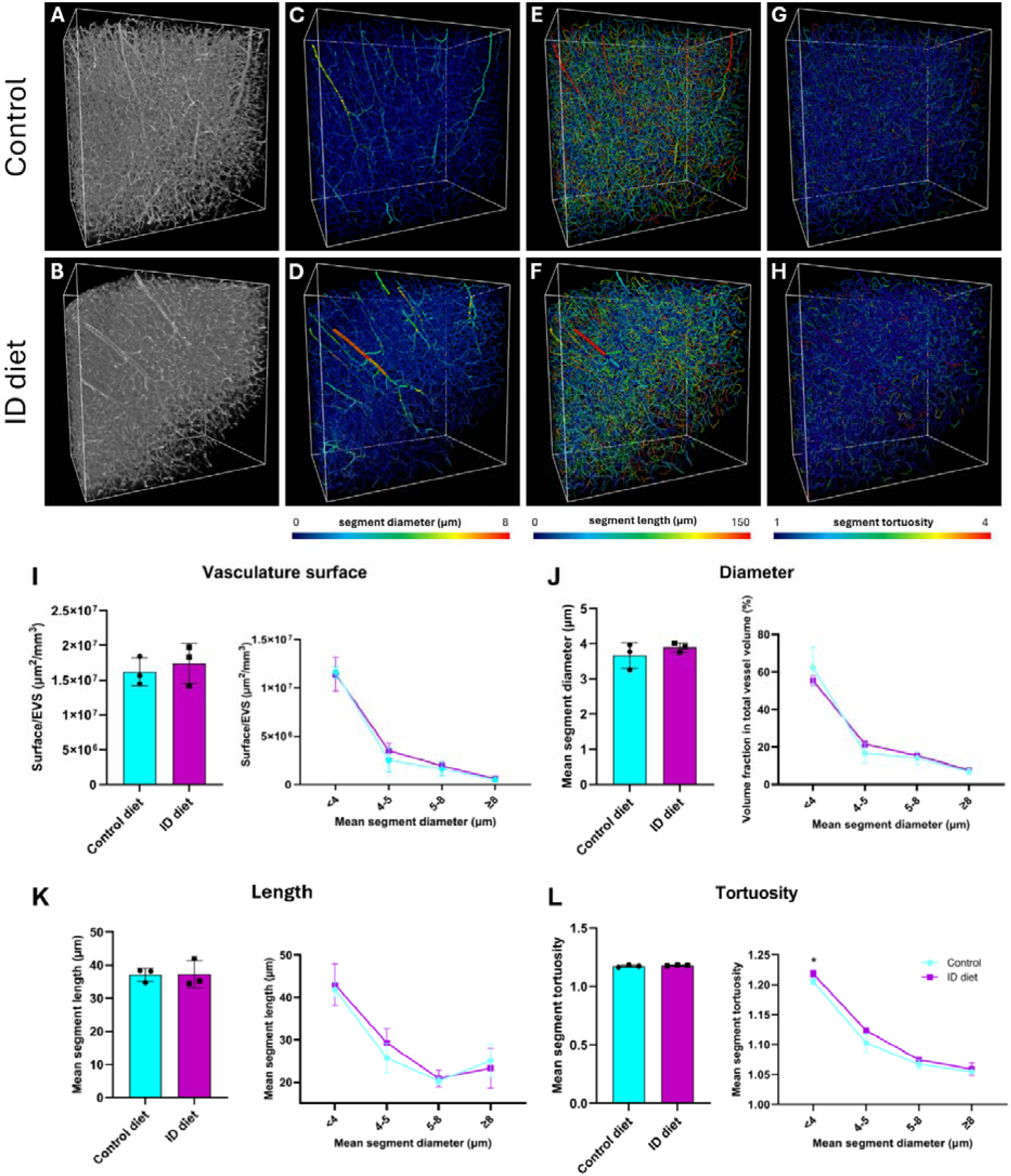
Analysis of effects of dietary ID on brain angioarchitecture. 3D reconstruction of preprocessed images of cortical vasculature labelled with intravenous injection of wheat germ agglutinin in control (**A**) and ID (**B**) mice. Spatial graphs of the vascular networks extracted from images in A and B representing the total network (barplots) and data binned by segment diameter (lineplots) with color coding according to the vessel segment diameter (**C, D**), length (**E, F**) and tortuosity (**G, H**). Data describing vascular network surface (**I**), vessel segment diameter (**J**), length (**K**) and tortuosity (**L**) with mice shown fed a control (turquoise) or ID (magenta) diet. In spite the tortuosity remained unchanged, a subpopulation reveals minor significant increase in the tortuosity of the small-caliber (<4 μm) vessels in ID. Data are analyzed using an unpaired t-test or Welch test (for tortuosity of subpopulations 5-8 μm and ≥8 μm). Data are depicted as mean ± SD for barplots and mean ± SEM for lineplots (n = 3 mice/group), *p < 0.05.

We then wanted to investigate the effect of ID on the content of TfR1 protein. Analyzing total brains and isolated brain capillaries, the content of TfR1 protein was seemingly identical both in total brain homogenates and in derivates of the isolated brain capillaries when comparing fractions of normal and ID mice (Fig. 5). The TfR1 level in control-fed mice ranged from 1.0 to 2.3 ng/µg protein in homogenates of total brain, and from 1.0 to 2.0 ng/µg protein in ID mice. In the derivates of the isolated brain capillaries, the capillary enriched fraction from control feed mice had a TfR1 content varying from 1 to 2.2 ng/µg protein, and from 1.1 to 2.5 in ID mice. The supernatant containing the post-vascular fraction varied in content from 2.5 to 6.4 ng/µg protein in control feed mice and from 2.9 to 6.8 ng/µg protein in ID mice (Fig. 5).

**Figure 5.**
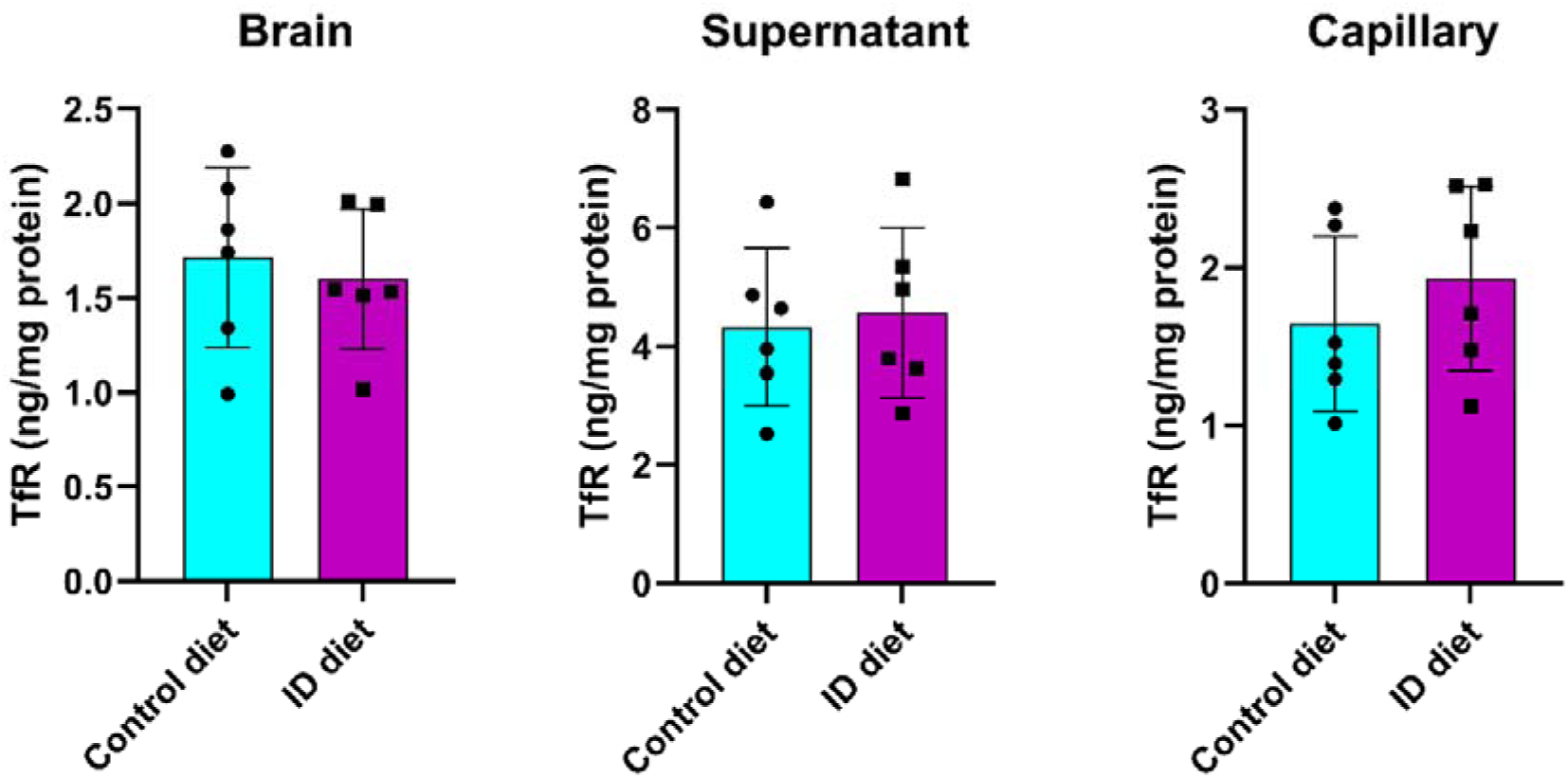
Transferrin receptor 1 (TfR on the Y-axis) protein in mice fed a control (turquoise) or ID (magenta) diet. Feeding the mice with the ID diet for eight weeks leads to an unchanged content of TfR1 in homogenates of the total brain and isolated fractions of the brain pellet (capillary) or the supernatant containing the post-vascular capillary depleted compartment of the brain. Data are analyzed using an unpaired t-test. Data are depicted as mean ± SD (n = 6 mice/group).

Next, we wanted to study, whether ID could be quantitatively reflected by increased uptake of nanogold-labeled anti-TfR1 antibodies. The uptake was higher in the total brain of ID fed mice increasing from approximately 104.1 ± 9.9 ng Au/µg protein to 139.1 ± 7.4 ng Au/µg protein (Fig, 6). The uptake was also higher in the post-vascular fraction (supernatant) of the capillary isolation increasing from 107 ± 10.9 to 140.3 ± 16 ng Au/µg protein, whereas the accumulation of nanogold-Ri727 in the capillary fraction remained unchanged when compared to that of the control fed mice (Fig. 6). Among extracerebral tissues, the uptake of Au-Ri727 was significantly higher in the liver (p < 0.005), spleen (p < 0.05), lung (p < 0.01), and retina (p < 0.0001) of mice fed the ID diet. The uptake in the liver was particularly high rising from 3501 ± 494.6 to 4581 ± 429.7 ng Au/µg protein and in the retina leading to an increase in uptake from approximately 9.5 ± 1.4 ng Au/µg protein to 12.6 ± 1.2 ng Au/µg protein (Fig. 6). Together these data showed that dietary ID lead to increased uptake of Au-Ri727 in tissues with prominent expression of transferrin receptors like brain, retina, spleen, and liver.

**Figure 6.**
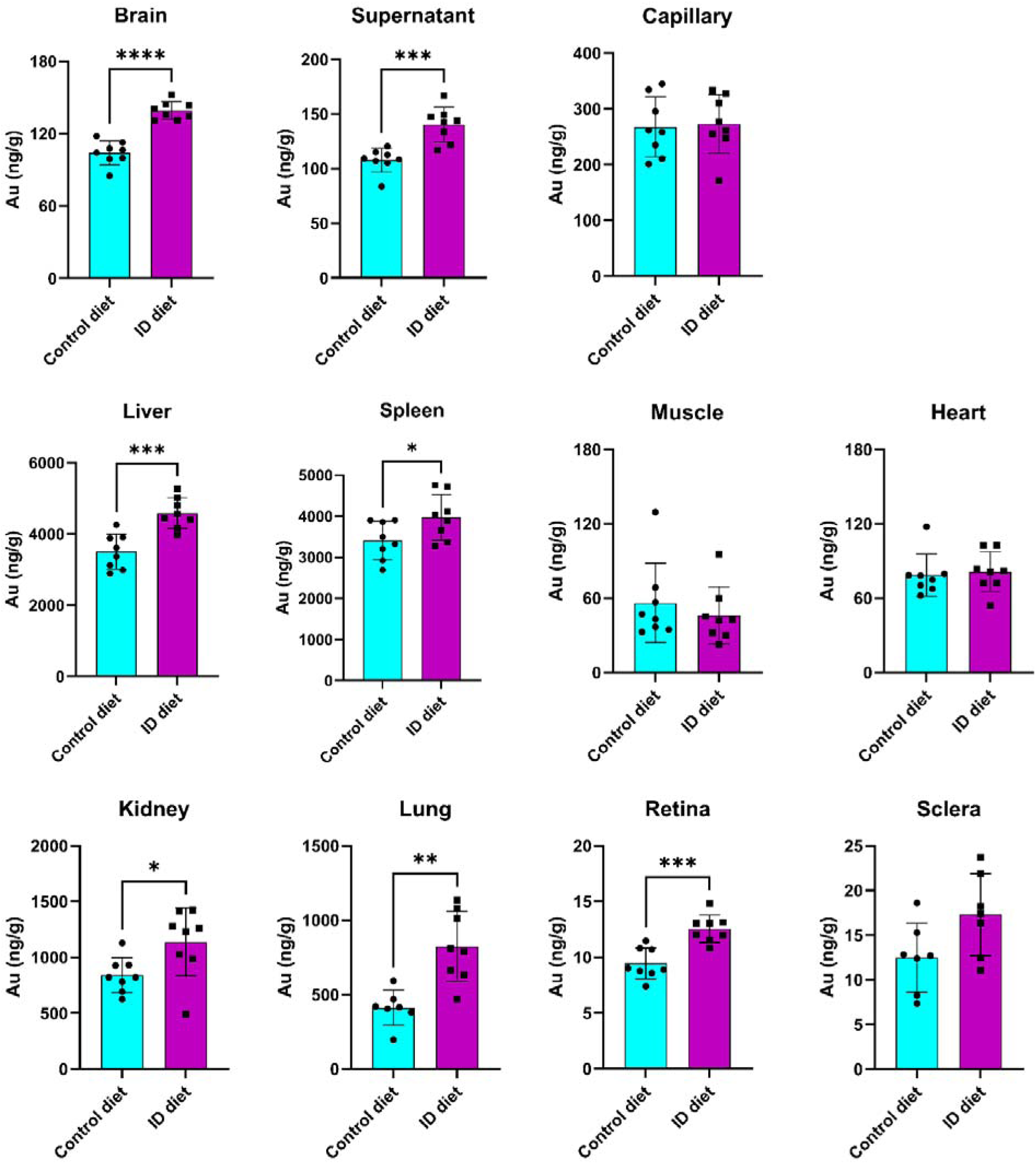
Measures of 1 nm nanogold (Au) conjugated to anti-TfR1 antibody in mice fed a control (turquoise) or ID (magenta) diet. Feeding the mice with the ID diet increases the Au-Ri727 uptake in the total brain (p < 0.0001) and the brain supernatant representing the post-vascular brain compartment (p < 0.001). The content of Au-Ri727 is not higher in the vascular fraction of the brain capillary isolation. Higher uptake of Au-Ri727 is seen in the liver (p < 0.001), spleen (p < 0.05), kidney (p < 0.05), lung (p < 0.01), and retina (p < 0.001). Data are analyzed using an unpaired t-test. Data are depicted as mean ± SD (n = 7-8 mice/group).

## Discussion

Subjecting adult mice to a diet with limited iron content for eight weeks resulted in ID documented by a significant lowering of iron content in several non-cerebral organs. The iron content in the brain remained unchanged compared to control-fed mice. Our data on cerebral iron following dietary ID corroborates those of a recent study performed in C57Bl6 mice, which revealed non-significant changes when subjecting adult female mice to changes in dietary iron for six weeks (Palsa et al., 2024). The preservation of iron in the brain is important as this shows that the brain was functioning with respect to iron uptake in spite of achieving peripheral ID. The content of copper was slightly changed in brain and skeletal muscles, but otherwise constant. The change in copper was expected, as dietary ID was previously shown to elevate copper in the brain (Monnot et al., 2011; Burkhart et al., 2024).

The iron content in isolated brain capillaries was lower in ID and supports the notion that cells in the brain can maintain iron during ID (Han et al., 2003). Expectedly dietary ID with lowering of peripheral tissue iron results in an elevated content of TfR1 to enhance the uptake of available iron (Teh et al., 2024). However, this was not observed in the present study as our data revealed that brain capillaries had similar content of TfR1 protein irrespective of iron status. Corroborating this observation, TfR1 levels relative to actin remained at a constant level in protein blots of female mice subjected to dietary ID (Palsa et al., 2024).

Studies in rodents have revealed that the uptake of iron-transferrin and expression of TfR1 at the BBB exerts plasticity in the brain in two particular conditions, i.e. during postnatal development and in ID. Age rather than iron status is decisive for TfR1 expression in brain capillaries (Moos et al., 1998; Moos and Morgan, 2001; Isasi et al. 2022). In rodents, the gestational period principally covers the first two trimesters in humans, with the transition between 1st and 2nd trimesters occurring only two days before delivery, whereas the third trimester in human pregnancy is adequate to the first weeks after birth (Markova et al., 2019). Concerning the early postnatal rat brain, a high expression of TfR1 leads to raised uptake of iron-containing transport in brain capillaries and subsequent higher transport of iron into the brain peaking at P15 postnatal days and rapidly declining when compared to the maturing brain at later ages (Taylor and Morgan, 1990). The neonatal brain represents a stage with particularly high expression of TfR1 (Moos et al., 1998) and histologically detectable iron, which disappears already around postnatal days 3-5 (Connor et al., 1995; Moos, 1995). The higher expression of TfR1 in the neonatal brain correlates with proliferation of the brain capillaries of the growing brain (Robertson et al., 1985) and compared to the maximal uptake of iron-transferrin, which occurs around postnatal days 10-15, this is not reflected in an adequately higher transport of iron or antibodies to the transferrin receptors across the BBB (Moos and Morgan, 2001), possibly because the BCECs at this early developmental stage are too undifferentiated to enable polarized vesicular trafficking.

Using iv injection of an anti-rat TfR1 antibody (OX26) for targeting BCECs in postnatal rats revealed a peak in the uptake around postnatal day 15, which was in good accordance with the notion of a developmentally higher uptake of transferrin at this age (Taylor and Morgan, 1990). In contrast, examining the effect of dietary ID on the uptake of OX26 in the developmental rat did not evoke a further higher uptake on P15, which likely could be attributed to that the normal rat at this developmental stage already has a stage of physiological ID, which means that the expression of TfR1 has reached a maximum and is unable to increase even further in spite ID (Moos and Morgan, 2001).

The developmentally regulated lower expression of TfR1 from postnatal ages older than P15 and onwards provided a more optimal opportunity to examine whether the effects of dietary ID would be able to raise TfR1 in BCECs. Our data indicated that TfR1 in BCECs remained constant despite systemic ID; an observation that was also observed prior at the morphological level in adult rats aged P70 subjected to dietary ID (Moos et al., 1998). The expression of TfR1 mRNA is post-translationally regulated by interactions of iron-responsive protein (IRE)s 1 and 2, which interact with an iron-responsive element (IRE) present in the 3’UTR end in conditions with depleted iron leading to stabilization of TfR1 mRNA with potential of higher mRNA translation (Rouault, 2013). The expression of TfR1 mRNA appears fairly constant with increasing age (Dornelles et al., 2010) and iron status (Han et al., 2003) and suggests that the lower content of TfR1 with increasing age is regulated by other mechanisms than available iron, which otherwise would have led to higher TfR1 in adult mice. It should not be overlooked that a recent study examining adult C57Bl6 mice with dietary ID found that males opposed to females had a higher content of TfR1 (Palsa et al., 2024). The sex difference was also reflected by the higher uptake of iron-transferrin from blood plasma in male mice with ID, but correspondingly to prior studies in rats, the uptake of iron-transferrin in adult mice was clearly exceeded when examining developing mice at P15 (Palsa et al., 2024).

The mechanisms leading to transport across the BBB can be considered a process of three successive steps, i.e. a) binding of the antibody to the TfR1 depending on the affinity, followed by receptor-mediated endocytosis of the TfR1-Ri7217 complex, b) transport of the TfR1-Ri7217 complex confined in vesicles through the BCECs to the abluminal side, and c) attachment and fusion of the transport vesicle with the abluminal membrane followed by a release of TfR1-Ri7217 into the brain interstitial space with recycling of the TfR1 receptor to the luminal membrane (Lee et al., 2000). Contradicting this notion, prior examination in the rat targeting the TfR1 in BCECs using a high-affinity anti-transferrin receptor IgG1 antibody (OX26) in adult rats revealed non-significant transport across the BBB as documented by clear tendency of the antibody to remain bound to TfR1-containing transport vesicles within BCECs and a minimum of transcytotic transport across the BBB (Moos and Morgan, 2001). A major question raised in the present study therefore concerned whether dietary ID would elevate the propensity of transport across the BBB of a TfR1-targeted antibody with high affinity for the receptor. Our data indicate that this was the case judging from the higher uptake of nanogold-conjugated Ri1727 in the total brain and the supernatant fraction of the capillary separation experiment containing the brain parenchyma of mice fed the ID diet.

An important question related to how Ri7217 can cross the BBB. A recent study implied that the affinity of antibodies crossing the BBB should be identified by means of the individual kinetic components ka (on-rate) and kd (off-rate) to characterize their KD (affinity constant) (Edavettal et al., 2022). This study demonstrates elevated transport across the BBB despite a high on-rate suggesting the off-rate is correspondingly raised, which might explain why the Ri7217 can pass the BBB. When attempting to explain why the transport of Ri7217 through BCECs is higher in ID, it should also be emphasized that non-vesicular transfer is significantly raised both with respect to albumin (Taylor et al., 1991) and non-immune IgG (Moos and Morgan, 2001). This suggests that a contribution of transendothelial trafficking through BCECs in ID comes from non-receptor mediated transport of Ri7217. BCECs express calcium (Ca^2+^)/calmodulin-activated kinase (CAMKK2) (De Bock et al., 2013) that controls TfR1-mediated trafficking (Sabbir, 2020), and it cannot be excluded that transport of TfR1-targeted vesicles through the BBB is raised in ID by changes in CAMKK2-CAMK4 signaling.

The current knowledge on the influence of ID on the microvasculature structure in the adult brain is limited. However, there is good evidence that iron is important for maintaining the structural integrity of the vessel wall, especially in the developing brain (Bastian et al., 2015; Simpson et al., 2015; Isasi et al., 2022), probably due to iron being co-factor for the iron-containing enzymes prolyl and lysyl hydroxylases both important for collagen IV synthesis (Commack et al. 1990). As the dietary ID regimen employed here decreased the iron content of the brain capillaries, which potentially could contribute to pathology, we investigated the state of microvascular architecture in the brain as a comprehensive, albeit indirect measure of vascular health. Existing reports have demonstrated an increase in retinal vasculature diameter and tortuosity in adult human subjects with ID (Shah and Modi, 2016). In neonatal rats, a compensatory increase in vessel density occurs in the hippocampus and even to a larger extent in the cerebral cortex following gestational ID (Bastian et al., 2015).

Given the importance of iron in the oxygen supply, maintenance of the vessel wall, and vascular tone, changes in angioarchitecture could theoretically also be expected in the present study. We, nevertheless, observed a largely preserved angioarchitecture. Similarly, we did not observe changes in vessel rarefication in dietary ID, which could have occurred due to malnutrition (Bastian et al., 2015), leading to changes in the ratio between the surface of the perfused endothelium and the extravascular space. These observations suggested that the vascular network in the current model of ID can provide proper perfusion of the associated parenchyma. The lack of vessel fraction change suggested that ID did not promote any compensatory angiogenesis, as there was no increase in the vessel fraction relative to the parenchyma. The only significant change we observed concerning the angioarchitecture in ID was an increase in vessel tortuosity. This change was nonetheless quite limited and significant only for the small-caliber vessels, meaning that despite indicating early stages of basement membrane disruption, further studies are needed to establish this unambiguously. In all, our model of dietary ID subjected to adult mice suggested preservation of the brain capillaries with unimpaired capacity for binding and transport of TfR1 targeted antibodies to a higher degree than seen in normal-fed mice.

## List of abbreviations (not complete)

BBB: blood-brain barrier;
BCEC: brain capillary endothelial cell;
ICP-MS: Inductively coupled plasma mass spectrometry;
ID: iron deficiency;
IDS: iduronate 2-sulfatase;
IV: intravenously;
MPS II: mucopolysaccharidosis type II;
Ri7217: high-affinity rat anti-mouse transferrin receptor IgG1 antibody;
SD: standard deviation;
SEM: standard error of the mean;
TfR1: transferrin receptor 1;
THF: tetrahydrofuran;
VOI: volume of interest;
WGA: Wheat Germ Agglutinin

## Declarations

## Acknowledgments

The 3D confocal imaging was performed using equipment at the Core Facility for Integrated microscopy, Department of Biomedical Sciences, University of Copenhagen. Steinunn Helgudóttir is acknowledged for performing ELISA analysis for TfR1.

## Ethics

The Danish Animal Experimental Inspectorate approved all experiments using experimental animals (License: no. 2013-15-2934-00893 and 2018-15-0201-01550).

## Consent to publish

The study does not include work on humans.

## Data availability

The datasets generated during and/or analyzed during the current study are available from the corresponding author upon reasonable request.

## Author contributions

Serhii Kostrikov (SK) and Kasper Bendix Johnsen (KBJ) designed and equally contributed to the study. Material preparation, data collection, and analysis were also performed by SK and KBJ. The first draft of the manuscript was written by SK and Torben Moos. Graphic illustrations were prepared by SK and Annette Burkhart. All authors commented on previous versions of the manuscript and approved the final manuscript.

